# High resolution imaging of nascent mitochondrial protein synthesis in cultured human cells

**DOI:** 10.1101/2020.05.05.076109

**Authors:** Matthew Zorkau, Christin A Albus, Rolando Berlinguer-Palmini, Zofia MA Chrzanowska-Lightowlers, Robert N. Lightowlers

## Abstract

Human mitochondria contain their own genome, mtDNA, that is expressed in the mitochondrial matrix. This genome encodes thirteen vital polypeptides that are components of the multi-subunit complexes that couple oxidative phosphorylation (OXPHOS). The inner mitochondrial membrane that houses these complexes comprises the inner boundary membrane that runs parallel to the outer membrane, infoldings that form the cristae membranes, and the cristae junctions that separate the two. It is in these cristae membranes that the OXPHOS complexes have been shown to reside in various species. The majority of the OXPHOS subunits are nuclear-encoded and must therefore be imported from the cytosol through the outer membrane at contact sites with the inner boundary membrane. As the mitochondrially-encoded components are also integral members of these complexes, where does nascent protein synthesis occur? Transcription, mRNA processing, maturation and at least part of the mitoribosome assembly process occur at the nucleoid and the spatially juxtaposed mitochondrial RNA granules, is protein synthesis also performed at the RNA granules close to these entities, or does it occur distal to these sites ? We have adapted a click chemistry based method, coupled with STED nanoscopy to address these questions. We report that in human cells in culture, within the limits of our methodology, the majority of mitochondrial protein synthesis occurs at the cristae membranes and is spatially separated from the sites of RNA processing and maturation.

## Introduction

Following the original observation of microcompartments within the mitochondrion (1, 2) researchers have been driven to investigate the organelle in ever increasing detail. We have known for many years that mitochondria contain two membranes. More recently, the microarchitecture of the organelle has been investigated following the establishment of fluorescent tools and high resolution light and electron microscopy or tomography (for reviews see (3-5)). It is now clear that the inner mitochondrial membrane (IMM) has highly defined regions. It comprises i) the inner boundary membrane (IBM) that is closely apposed to the outer mitochondrial membrane (OMM) and where contact sites containing the machinery for importing proteins from the cytosol are found (6, 7), ii) highly invaginated cristae membranes (CM) that house the majority of the OXPHOS complexes (8-10), in particular the FoF1 ATP synthase, dimers of which help generate the architecture of the membranes (11) and iii) large (10 – 40nm) complexes termed cristae junctions (CJ), composed of the *M*itochondrial contact site and *C*ristae *O*rganising *S*ystem (MICOS; (12-14)), which forms the cristae and separates them from the IBM.

There has also been much focus on the mitochondrial genome, how and where it is expressed. This has led to the identity and characterisation of the nucleoid (15-18) and more recently the mitochondrial RNA granule, a complex formed by phase transition where transcripts are processed and matured (19-21). Translation occurs on membrane-associated mitochondrial ribosomes, which are also at least partially assembled in the RNA granule (22, 23). Exactly where, however, protein synthesis is performed in human mitochondria, whether at the CM, CJ or IBM, within, close to, or distal to the RNA granules, is unknown. To follow this process in yeast mitochondria, Jakobs and colleagues harnessed an elegant series of experimental approaches (24). The yeast *S. cerevisiae* express a battery of translational activators that show specificity to mt-mRNAs, which encode individual components of OXPHOS complexes III, IV and V (25). Using immuno-labelling super resolution and electron microscopy, data were produced that were consistent with mtDNA-encoded complex V subunits being synthesised predominantly on the cristae membranes, whilst complex III and IV components were synthesised both at the IBM and CM. This approach is not feasible in human cells as with the exception of TACO1 (26), human mitochondria do not contain translational activators, moreover the majority of mtDNA encoded human proteins are components of complex I, an OXPHOS complex that is absent in *S. cerevisiae*. As translational activators were used as surrogate markers for translation, a direct assay for protein synthesis would now be ideal. One study has utilised a pulse-labelling click chemistry method to directly measure mitochondrial protein synthesis in intact human cells (27). However, the limited resolution and depth of analysis meant many spatial characteristics of newly synthesised mitochondrial proteins remained undefined. In particular, no information of the sub-mitochondrial localisation of protein synthesis in human cells could be obtained.

We report here a comprehensive set of analyses that directly measure spatiotemporal kinetics of mitochondrial protein synthesis. Click chemistry (28) is the technique of choice for this approach, appealing to cell biologists as it utilises free azide or alkyne labelling moieties that are rarely found in cells. The non-canonical methionine analogs homopropargylglycine (HPG) or azidohomoalanine (AHA) can be substituted for methionine and subsequently visualised in fixed cells with azido or alkyne fluorophores respectively, in a process referred to as fluorescent non-canonical amino acid tagging, or FUNCAT (29, 30). We have adapted initial protocols (27, 31) and used HPG whilst inhibiting cytosolic protein synthesis to specifically visualise mitochondrial protein synthesis with both confocal microscopy and super resolution STED nanoscopy. Signals reporting protein synthesis can be detected in various human cell lines after 5 minutes. Over 90 minutes of HPG labelling, the proportion of translationally active mitochondrial network (∼50%) remained relatively unchanged. Using this method, we can report that the majority of protein synthesis occurs at the cristae membranes. Further, our STED nanoscopy revealed that synthesis initiates mainly at sites separated from RNA granules, suggesting either that if the mitoribosome is loaded with mt-mRNA close to the RNA granule it must be able to travel to the cristae membrane prior to synthesis, or that the mitoribosome is loaded at the cristae membrane itself.

## Results

### Measuring spatiotemporal kinetics of mitochondrial protein synthesis in human cells

Originally, a method had been described to measure newly synthesised cytosolic proteins in intact cells by immunofluorescence. This involved cells in methionine-free media being incubated with the alkyne or azido-methionine derivatives, HPG or AHA, respectively. These methionine analogs were incorporated into nascent protein, with visualisation occurring after fixation and copper-catalysed cycloaddition of the azido or alkyne fluorophore (29, 32). We adapted this method, termed FUNCAT (fluorescent non-canonical amino acid tagging), to exclusively measure mitochondrial protein synthesis and further refined previous techniques (Fig. S1 (27, 31)). To ensure the method could be widely applied and that observations were robust, all experiments were performed with at least three types of human cultured cells, unless otherwise indicated. The cell lines included two transformed lines (HeLa and U2OS) and primary dermal fibroblasts. As can be seen in Fig. 1A (U2OS cells) and *SI Appendix* Fig. S1 (HeLa, fibroblasts), co-treatment of cells with cycloheximide (to inhibit cytosolic protein synthesis) and HPG allowed visualisation of a signal that colocalised with the mitochondrial network. The HPG signal was lost upon simultaneous inhibition of cytosolic and mitochondrial protein synthesis (Fig. 1B), demonstrating that the signal in Fig. 1A exclusively represented HPG-labelled mitochondrially synthesised proteins. Although this demonstrated detectable mitochondrial translation, it did not determine whether HPGylated mt-tRNA^Met^ could be used to initiate synthesis. To address whether the signal represented only elongation of previously initiated proteins, cells were pretreated with puromycin to terminate synthesis, prior to addition of HPG. As shown in *SI Appendix* Fig. S2, mitochondrially-derived signal was observed after puromycin pretreatment, consistent with *de novo* initiation of translation. This indicated that HPGylated mt-tRNA^Met^ could initiate synthesis as well as elongate nascent peptides. It is tempting to infer that a subset of HPGylated mt-tRNA^Met^ could have been formylated by mitochondrial methionyl-tRNA formyltransferase, mimicking the endogenous initiation mechanism (33), but recent data suggests under certain circumstances initiation in mitochondria may occur without formylation of an initiating mt-tRNA^Met^ (34).

**Figure 1.**
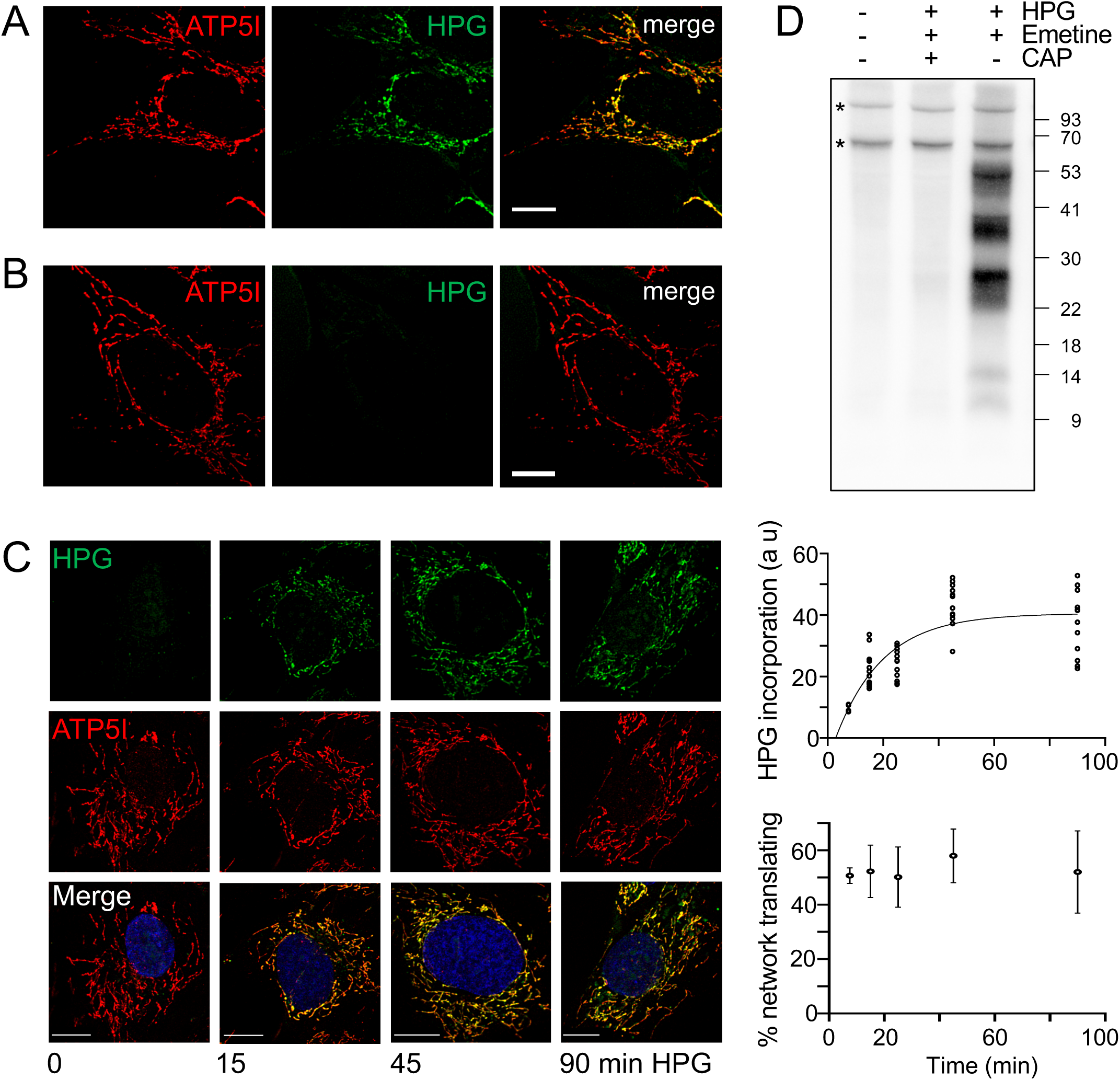
Translation of mtDNA encoded proteins can be monitored by incorporation of HPG in a time dependent manner revealing the distribution across the mitochondrial network. U2OS cells were pulsed with HPG (25 min) and cycloheximide in the absence (A) or presence (B) of chloramphenicol. Cells were then fixed, underwent the click reaction and were stained with antibodies against ATP5I. (C) Control fibroblasts were cultured in HPG with inhibition of cytosolic translation by cycloheximide for the time periods indicated. Post-fixation, click reactions were performed and immunostained as for A and B. All images are representative, scale bars = 10 μm. The total HPG incorporation as a marker of mt-protein synthesis was plotted over a 90 minute time course (upper graph). The proportion of the mitochondrial network stained with HPG reflecting how much of the reticulum is translationally active was calculated as a percentage from multiple images (n=4, 7.5 min; n=12, 15, 25, 45, 90 min) from each time point and presented graphically (lower graph). All experiments were performed as a minimum of 3 independent experiments. (D) Wild type HEK293 cells were incubated in the presence or absence of inhibitors followed by addition of HPG. Mitochondria were isolated and the click reaction performed as described in Methods. Samples were separated by 12% Bis-Tris PAGE and biotinylated proteins visualised with Streptavidin-HRP and ECL. * endogenously biotinylated mitochondrial proteins.

To determine whether the mitochondrial FUNCAT assay could be utilised to characterise the location and rates of mitochondrial translation, HPG incorporation in the mitochondrial network were measured over a time course (Fig. 1C). The rate in fibroblasts was relatively linear over a 45 minute pulse but a clear plateau was evident by 90 minutes (Fig. 1C, upper graph), which reflected trends seen in ^35^S-met radiolabelling of mitochondrial protein synthesis after similar inhibition of cytosolic protein synthesis.

It is not currently known whether translation occurs in discrete foci or is evenly distributed throughout the mitochondrial network. Analysis of fibroblasts revealed that after a 15 minute pulse a reasonably homogenous pattern of synthesis was present across the network (Fig. 1C, lower graph). Longer pulses in these and HeLa cells reflected only an increase in intensity rather than redistribution of signal (Fig. 1 and *SI Appendix* Fig. S3). More detailed analysis revealed protein synthesis to be modestly but significantly enriched in all three cell lines tested in perinuclear compared to peripheral mitochondria (*SI Appendix* Fig. S4).

The established method for visualising human mitochondrial protein synthesis requires ^35^S-met metabolic labelling (35) and has been adopted by many mitochondrial laboratories over the past 20 years. We attempted to visualise the newly synthesised HPGylated proteins in gel based systems to avoid the dependency on radiolabel. Following HPG pulse labeling, mitochondria were isolated and HPGylated proteins labelled with picolyl biotin permitting visualisation after transfer. Robust incorporation of HPG into the majority of mitochondrial proteins can be seen (Fig. 1D), with a band pattern resembling previous reports for ^35^S-met gels. Labelling was eliminated by chloramphenicol treatment, confirming that the signal represented mitochondrially encoded proteins. However, despite trialling many combinations, our optimised protocol and gel systems were unable to separate and identify all of the 13 species (*SI Appendix* Fig. S5).

### Turnover of newly synthesised HPGylated mitochondrial proteins

Although it is not immediately relevant to the question of where nascent synthesis is occurring in the mitochondrion, we were interested to know whether polypeptides containing multiple HPG molecules were being stably incorporated into OXPHOS complexes. A recent report suggested that in HeLa cells fully assembled complexes are highly stable, however, the majority of subunits are synthesised in substantial excess and the unassembled components are generally degraded within 3 hours (36). We therefore, elected to perform a pulse-chase experiment over 36 hours (Fig. 2A). Following HPG treatment (2 hours), both HPG and the reversible cytosolic protein synthesis inhibitor cycloheximide were removed and methionine added for the chase periods indicated. After 3 hours post HPG pulse, approximately 87 ± 15% of the signal remained suggesting that the majority of newly synthesised proteins were not rapidly degraded. Although there was a subsequent decrease after 6 hours (36 ± 16%), inferring that a proportion of the newly synthesised mitochondrial protein is not assembled, retention of signal at 36 hours confirmed incorporation of at least a subset of labelled protein into stable complexes.

**Figure 2.**
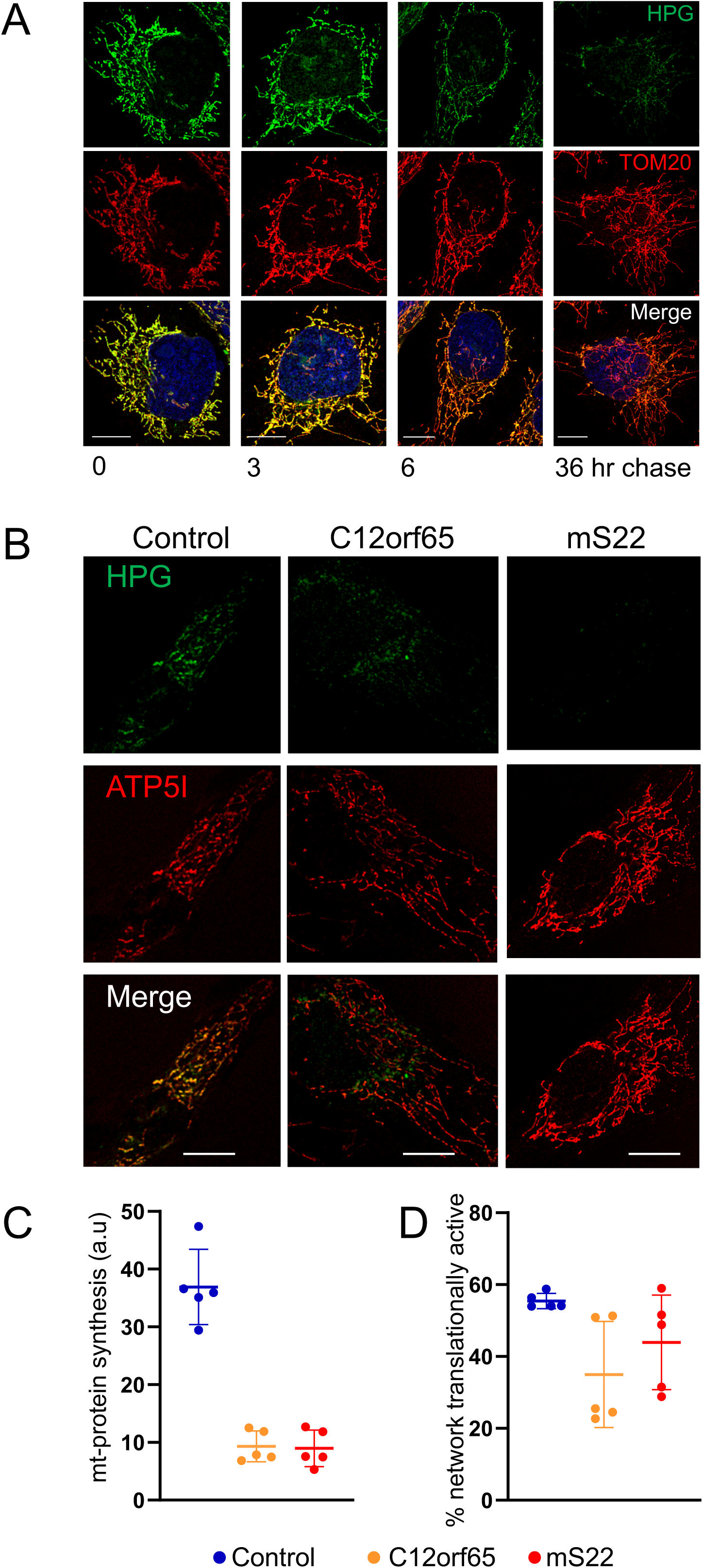
HPG remains detectable in mitochondrial networks after 36 hours, and incorporation can reflect translation defects in mitochondrial patient cell lines. (A) U2OS cells were pulsed with HPG (green) for 2 hours followed by increasing chase length as indicated. Click reactions were performed as previously described. Mitochondrial outer membranes were visualised with antibodies against TOM20 (red) and nuclei were stained with Hoechst (blue). Representative deconvolved confocal images are shown, all scale bars = 10 μm. (B) Dermal fibroblasts from patients with mutations in nuclear genes (C12orf65 and mS22) known to cause defects in mitochondrial translation were pulsed with HPG (45 min) alongside a control line. The mitochondrial network was visualised with antibodies against ATP5I. Representative deconvolved confocal images are shown, all scale bars = 10 μm. (C) Mitochondrial protein synthesis (HPG signal normalised to mitochondrial area; arbitrary units) was quantified for each cell line, as was the proportion (%) of the mitochondrial network where translation was occurring (D). Each data point (n=5) represents analysis within an individual cell.

### Mitochondrial FUNCAT can be used to identify mitochondrial protein synthesis defects

Traditionally, to determine whether individuals have defects in mitochondrial protein synthesis, patient derived cell lines have been subjected to metabolic labeling with ^35^S-met in the presence of a cytosolic protein synthesis inhibitor (35). One of our goals was to establish mitochondrial FUNCAT as a tool that overcame the need for radiolabel in such assays. To establish whether our methodology could be applied to identify such defects, we used our assay to measure mitochondrial protein synthesis normalised against mitochondrial unit area in control fibroblast and two patient cell lines. Both patients carried defects in mitochondrial protein synthesis – a homozygous exonic deletion (c.210delA, p.(Gly72Alafs ^*^13) frameshift mutation in the release factor C12orf65 (37) and a mutation in the mitoribosomal subunit mS22. In both cases, a substantial decrease in signal was noted after 45 minutes when compared to controls (Fig. 2B, C), but there was no marked variation in the proportion of translationally active network (Fig. 2D). Importantly, by 90 minutes, the difference in signal between patient and control line was abolished (*SI Appendix* Fig. S6). This emphasised that kinetics have to be carefully established for each cell line. These data confirm that the mitochondrial FUNCAT assay is suitable for identifying and measuring mitochondrial protein synthesis defects and potentially any changes in distribution of signal within the network.

### Nascent protein synthesis occurs predominantly at the cristae membranes

The main focus of this study was to establish the sub-mitochondrial localisation of nascent protein synthesis. To achieve this goal we adapted the mitochondrial FUNCAT technique to enable visualisation by super resolution STED nanoscopy. One strength of FUNCAT is that fluorophores are attached directly to the methionine analog by the copper catalysed alkyne-azide reaction. We were able to optimise the technique using the picolyl Alexafluor 594 azide. Colocalisation was then determined with various markers using immunofluorescence techniques as described. We first assessed whether newly synthesised proteins could be identified away from the mitochondrial outer membrane (Fig. 3A). HPG was pulsed for 30 minutes and the signal was compared to that produced from antibodies specific for the outer membrane marker TOM20. High resolution imaging indicated that new proteins are synthesised at a location internal to the outer membrane (Fig. 3A, B). Further, the HPG signal clearly shared a location with ATP5I, a component of the FoF1 ATP synthase and member of the CM (Fig. 3B, C). Therefore, to assess more precisely where in the inner mitochondrial membrane the highly hydrophobic mtDNA encoded proteins were being inserted, we quantified colocalisation with markers from the inner boundary membrane (IBM; TIM23 a component of the inner membrane translocase), the cristae junctions (CJ; mic60, a member of the MICOS complex) and a second marker of the CM cristae membrane (COXI, a component of cytochrome *c* oxidase or complex IV). Images in Fig. 3D confirm that the majority of HPG signal is found in the cristae membrane, colocalising with COX1 and only rarely in the IBM. MIC60, was used to visualise the cristae junction (38), where the IBM begins to invaginate to form the distinct compartment of the cristae membrane (Fig. 3D and *SI Appendix* Fig. S7 ATP5l *cf*. MIC60). Again, the majority of the HPG signal is distinct from and internal to MIC60 puncta, consistent with the newly synthesised proteins being inserted into the cristae membranes. Quantification using Manders’ colocalisation coefficient indicated that of the HPG signal 50.8 ± 8.8% associated with the cristae (COX1), whilst only 21.5 ± 8.9% colocalised with the IBM marker TIM23 and 12.6 ±. 5.4% with CJ marker MIC60. Analysis with Spearman’s rank correlation coefficient also revealed a 3-fold increase in correlative relationship between HPG and Cox1 or ATP5l signals compared to markers for the other sub-membrane compartments, a phenomenon also observed over longer pulses. Similar labelling is also seen when comparing to MIC10, a second member of the MICOS complex and marker of the CJ (*SI Appendix* Fig. S7). To account for the possibility of movement of newly synthesised proteins within the 30 minute pulse, a shorter (7.5 minute) pulse was applied that revealed a similar distribution (*SI Appendix* Fig. S3). However, the majority of the data analysed was after a 15 or 30 minute pulse, when the capacity to attain sufficient signal and signal:noise were significantly improved. A previous study using immuno-EM reported an enrichment of a single yeast mitoribosomal subunit YmL36 at the cristae membranes (9), an observation consistent with our STED studies with antibodies against uS15m (Fig. 4), which, crucially, visualised nascent rather than steady state distribution. Taken together, our data support the majority of protein synthesis occurring at cristae membranes.

**Figure 3.**
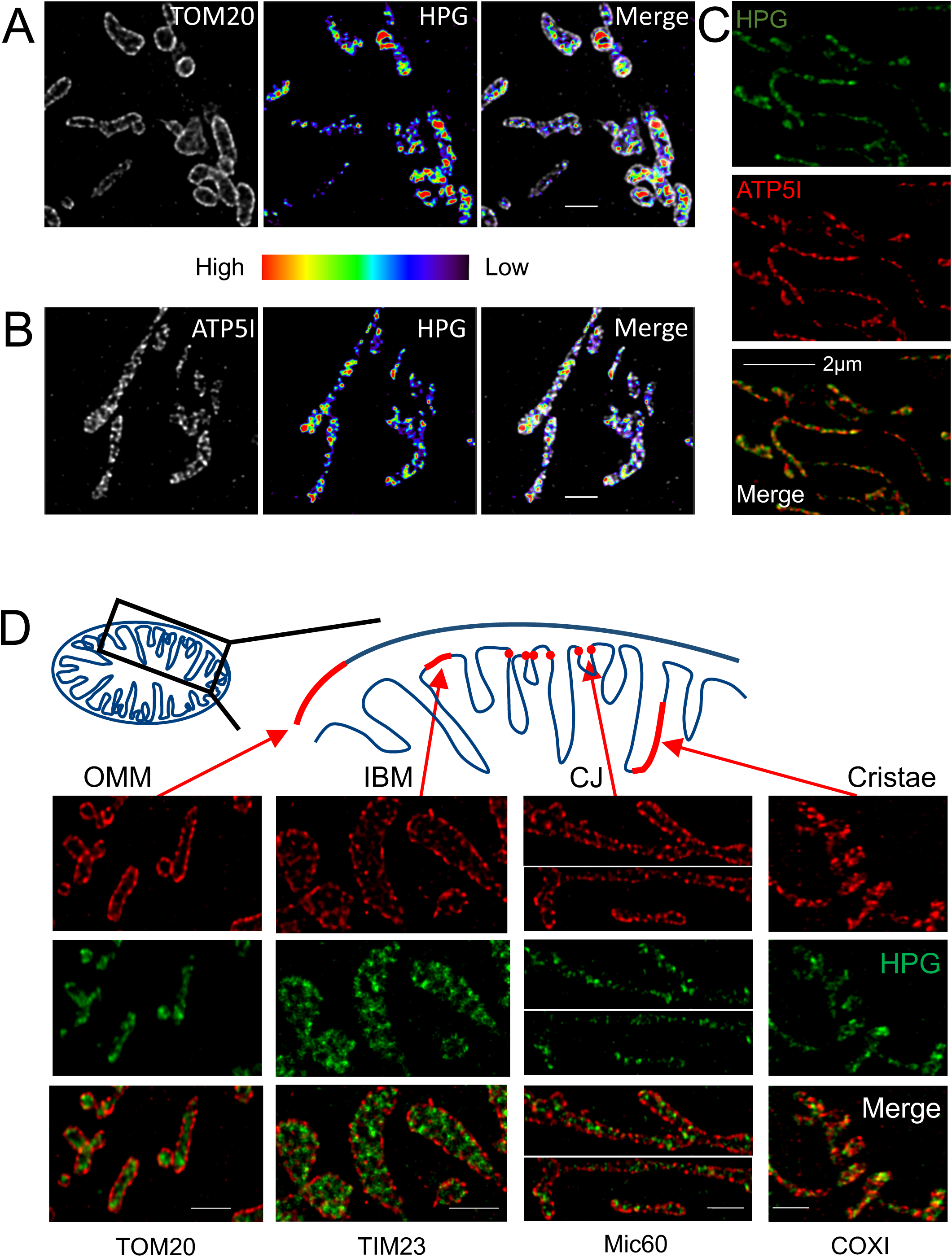
Intra-mitochondrial protein synthesis is enriched on cristae membranes. *De novo* protein synthesis was visualised in HeLa cell mitochondria through incubation with HPG (30 min) in the presence of cycloheximide. Cells were stained with antibodies to highlight the positions of the outer mitochondrial membranes (TOM20, panel A) or the cristae (ATP5I, panel B). The colour indicates relative enrichment of HPG incorporation (heat map range 3000 −11000) reflecting mt-translation. Representative deconvolved STED images are shown, with scale bars = 1 μm. (C) Control fibroblasts were incubated with HPG (25 min) in the presence of cycloheximide and cells stained with antibodies against ATP5I. Scale bar 2 μm. (D) A schematic depicting in red regions of the mitochondrial outer (OMM), inner boundary (IBM), cristae junction (CJ) and cristae membranes links to relevant panels below of U2OS cells pulsed with HPG (30 min) in the presence of cycloheximide co-stained with membrane specific antibodies and visualised with STED microscopy. MIC60 image split to remove central blank region. All scale bars are 1μm.

**Figure 4.**
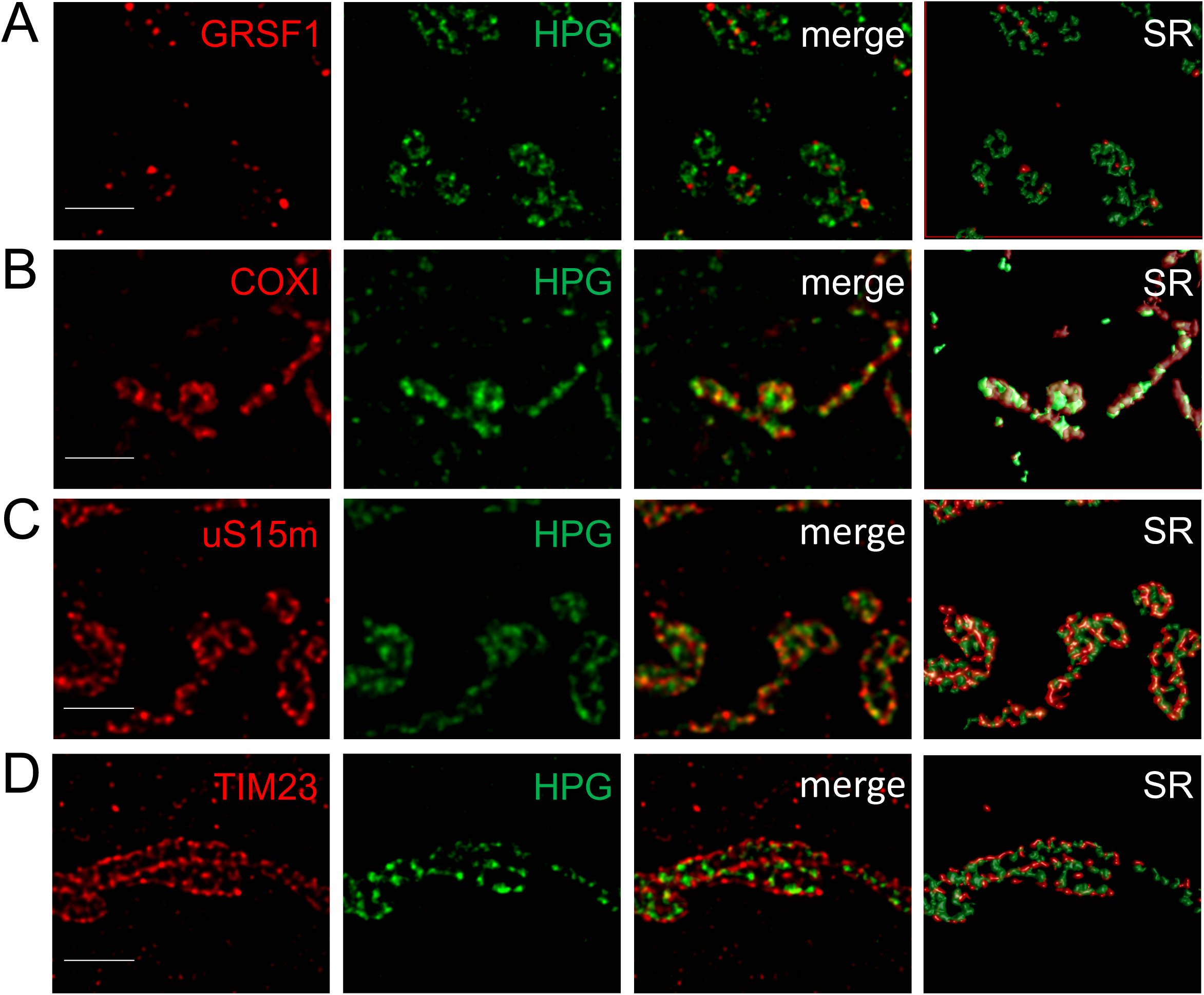
Spatial distribution of mitochondrial RNA granules and intraorganellar translation. U2OS cells were pulsed with HPG (15 min) prior to click reactions and fixation as per Methods. Immunostaining was performed with antibodies against markers of the mitochondrial RNA granules (GRSF1, panel A), cristae (COXI, panel B), a mitoribosomal protein of the small subunit (uS15m, panel C) or inner boundary membrane (TIM23, panel D). Representative deconvolved STED microscopy images are shown, as well as surface rendered (SR) representations of the merged data. All scale bars = 1 μm.

### Protein synthesis occurs at sites distal to the mitochondrial RNA granule

Mitochondrial gene expression appears highly compartmentalised, with large, juxtaposed complexes (the nucleoid and the RNA granule) in the mitochondrial matrix as highlighted in the introduction. Is it possible that translation also occurs within the mitochondrial RNA granule, or even in a third juxtaposed structure creating a form of production line to synthesise new proteins ? In yeast mitochondria, this idea has been supported by co-immunoprecipitation, proteomics and super resolution microscopy studies. These data revealed an association between the mitoribosomes and other gene expression proteins, leading to the concept of MIOREX, or *MI*tochondrial *OR*ganization of gene *EX*pression (39). To assess whether human mitochondria show a similar arrangement of associated complexes, we used STED nanoscopy to visualise nascent mt-protein synthesis (HPG) and antibodies to highlight two separate markers of the mitochondrial RNA granule (GRSF1, Fig. 4 and FASTKD2, *SI Appendix* Fig. S8). Figure 4 highlights that the majority of HPG signal is spatially distinct from mitochondrial RNA granules. Quantification (Manders’ colocalisation analysis) of data from the 15 minute pulse-labelled cells revealed that 64.4 ± 8.2% of the HPG signal colocalised with the mitoribosomal subunit uS15m but only 9.5 ± 2.9% with the RNA granule (GRSF1). Furthermore, at 30 minutes an approximately 8-fold enrichment of HPG signal was identified with the cristae marker ATP5l compared to GRSF1 (54.6 ± 1.4% *cf* 6.9 ± 2.1%). Even as early as a 7.5 minute pulse, the staining profile of HPG more closely resembled ATP5l than FASTKD2 (*SI Appendix* Fig. S8). The pattern of staining, assessed using surface rendered 3D maps, revealed that frequently but not exclusively, large stretches of HPG signal are adjacent to RNA granules. However, little evidence was found for compartmentalisation of protein synthesis within RNA granules.

## Discussion

We report that mitochondrial FUNCAT is a robust method for visualising protein synthesis in human cells. This technique represents a significant development in methodology to investigate human mitochondrial protein synthesis, in particular with an adaptation to higher resolution confocal and STED microscopy for the first time. Unlike ^35^S-met radiolabeling that was limited to scintillation counting or gel based display formats, this approach can visualise spatial information on the distribution of translation within the mitochondrial network. When implemented with super resolution microscopy and combined with immunofluorescence, as we have here, it allows us to gain a better understanding of how and when different machineries involved in all aspects of gene expression are integrated and how this is affected by different disease conditions. Currently, it is not possible for us to confidently produce consistent data with HPG pulse times shorter than ten minutes. For this reason we have exercised caution in our interpretations and conclusions. We believe our data is consistent with the majority of nascent protein synthesis occurring at the cristae membrane and distal from the nucleoid or RNA granule. One alternative explanation of our data could be that synthesis occurs at the inner boundary membrane close to the RNA granule and the new peptides are rapidly moved individually or as partially assembled complexes through the cristae junctions and into the cristae membranes. This is unlikely, as we see no major variation in the distribution irrespective of how long a pulse we use (15 - 120 minutes), merely an increase in the intensity of the signal. Second, calculation of diffusion coefficients in the cristae membrane show, for at least some members of the mature OXPHOS supercomplexes that reside in the cristae membranes, their diffusion rates are slow (10), reflecting the high protein content in these membranes that make rapid movement away from the site of synthesis improbable. Third, it has been argued that the CJs make a natural diffusion barrier in the inner membrane, effectively separating the IBM from the CMs (40, 41). Indeed, recent high resolution studies of live cells have shown that cristae can be biophysically distinct to the IBM as a consequence of CJs providing a form of physical insulation (42). Therefore, it is unclear how easy it would be for complexes made in the inner boundary membranes to diffuse through the constricted membranes around the MICOS complexes at the cristae junction. Our data consistently showed newly synthesised proteins occurring internal to OM, IBM and CJs.

At first sight, our conclusion may seem to contrast with the previous work by Jakobs and colleagues in yeast, which inferred that mitochondrial subunits of complexes III and IV are made at both inner boundary and cristae membranes dependent on their assembly status, whilst complex V is preferentially synthesised on cristae membranes (24). However, unlike that investigation that monitored mt-translation via mRNA-specific translational activators and assembly factors, mitochondrial FUNCAT gives a readout of all mitochondrial protein synthesis and cannot distinguish between components of different OXPHOS complexes. It is, therefore, possible that as with yeast complex V, human complex I components are largely made at the cristae membrane and as these constitute the major part of the mitochondrially-encoded proteome this may bias the data, whilst translation of the Complex III, IV and V components may be more widely distributed across cristae and IB membranes. Irrespective, it is clear that the main signal is visible first at the cristae membrane.

A second interesting observation was that synthesis did not appear to be at discrete foci within or apposed to the RNA granule, unlike the situation in yeast (39). Consistent with our observation is previous data based on quantitative immunoelectron microscopy that reported the majority of the yeast mitoribosomal subunit YmL36 to be associated with the cristae membrane and notably, when treated with puromycin to terminate synthesis, this selective localisation was lost (9). This may also be consistent with further work of Bogenhagen and colleagues (43) showing assembly of the human mitoribosome initiates close to the nucleoid or RNA granule but then has late assembled components potentially added at distal sites. Finally, another current limitation is that mitochondrial FUNCAT cannot allow synthesis to be followed in real time. Considering the superb nanoscopy images that are being generated to allow us to visualise the ultrastructure of the mitochondrion in real time (4, 42, 44, 45), we hope to soon adapt our protocol to follow mitochondrial protein synthesis in live cells.

## Materials and Methods

### FUNCAT - Cell culture and click chemistry labelling of mitochondrial translation in human cell lines

Cultured dermal human fibroblasts, U2OS or HeLa cells were grown in Dulbecco’s modified Eagle’s medium, (Sigma D6429) supplemented with 10% foetal calf serum (Sigma), 1x non-essential amino acids and 50 μg/ml uridine (Thermo Fisher Scientific) at 37°C in humidified 5% CO_2_. Access to samples that were excess to diagnostic requirements and were approved for research was covered by the license “Role of mitochondrial abnormalities in disease” (REC ref 2002/205) issued by Newcastle and North Tyneside Local Research Ethics Committee. For FUNCAT experiments, cells were first seeded onto glass coverslips and cultured for 1-2 days. *In vivo* labelling of mitochondrial translation products was performed by pulsing cells with methionine-free DMEM containing the methionine analogue HPG (Jena Bioscience) while cytosolic translation was inhibited with 50 μg/ml cycloheximide. When inhibiting mitochondrial translation, 100 μg/ml chloramphenicol was also added during the pulse period. If a chase was performed, methionine-free DMEM was replaced with full medium. Cells were pre-permeabilised with 0.005% digitonin in mitochondria-protective buffer (MPB: 10 mM HEPES/KOH, 10 mM NaCl, 5 mM MgCl_2_ and 300 mM sucrose in H_2_0, pH 7.5) for 80 secs at room temperature to remove unincorporated HPG before being fixed in pre-warmed 8% formaldehyde in MPB for 7 min. Cells were fully permeabilised with 0.5% (v/v) Triton X-100 in PBS (137 mM NaCl, 2.68 mM KCl and 10 mM Na_2_HPO_4_, pH 7.4) and subsequently blocked with 5% (w/v) BSA for 10 min. Pulse-labelled mitochondrial proteins were detected with a copper-catalysed azide–alkyne cycloaddition (600 uM copper sulphate, 1.2 mM BTTAA, 40 μM picolyl Alexa Fluor 555 (CLK-091-1) or 594 (CLK-1296-1) azide and 2 mM sodium ascorbate in PBS, all Jena Bioscience) click chemistry reaction for 40 min. Immunocytochemistry was used to co-label specific mitochondrial targets of interest. This involved incubation with a primary antibody diluted in 5% BSA for 1 hour at room temperature, followed by species-specific Alexa Fluor 532 (Thermo Fisher A-11009) or ATTO647N (Sigma M8645) secondary antibodies diluted 1:200 in 5% BSA for 40 min. Nuclei were stained with Hoechst for 5 min and cells were mounted with ProLong Glass Antifade mountant. Primary antibody details are listed in *S1 Appendix*, Table S1.

### Confocal imaging and 3D STED nanoscopy

Confocal imaging and 3D STED nanoscopy were performed on a Leica TCS SP8 gSTED 3X (Leica Microsystems) microscope equipped with white light lasers, HC PL APO 100x/1.40 Oil STED WHITE objective and 63x/1.40 Oil HC Pl Apo CS2 objective for confocal imaging were used. A voxel size of (35-40) x (35-40) x 130 nm (xyz) for x63 confocal and (10-20) x (10-20) x 100 nm (xyz)-nm for STED images was used. The fluorophore Alexa Fluor 532 was excited at 527 nm and STED was performed at 660 nm for super resolution imaging. Alexa Fluor 555 was excited at 555 nm and STED was performed at 660 nm. Alexa Fluor 594 was excited at 590 nm and ATTO647N at 646 nm, while STED was performed at 775 nm for both fluorophores. Images were deconvolved and, where indicated, rendered into computer-generated 3D surface maps, using Huygens software (Scientific Volume Imaging, WWW.svi.nl).

### Image analysis

In each case, unless otherwise stated, 5-12 images were analysed from a single representative experiment that was repeated at least three times with similar results. For determining amount and distribution of mitochondrial protein synthesis within whole cell confocal images, Columbus™ software (PerkinElmer) was used on raw images. A mitochondrial translation coefficient was determined by multiplying HPG surface area by mean fluorescence intensity and this was normalised to mitochondrial surface area (defined by labelling of a general mitochondrial marker such as ATP5I) to give mitochondrial protein synthesis per mitochondrial area. For characterising other spatial parameters or analysing STED data, deconvolution and pixel based colocalisation was applied (Scientific Volume Imaging). The colocalisation analysis utilised Spearman’s rank correlation coefficient to determine correlation between fluorophore intensities, while Manders’ colocalisation coefficient (M1 and M2) was used to calculate proportions of co-occurrence between 0 and 1.

### BONCAT - Cell culture and click chemistry labelling of mitochondrial translation in human cell lines

Wild type Hek293 cells were grown in DMEM (Sigma D6429) supplemented with 10% FCS, 1x non-essential amino acids and 50 μg/ml uridine (37°C, 5% humidified CO_2_). Medium was refreshed one day prior to the experiment and cells harvested at 50% - 80% confluency. Cells were depleted of methionine (Met) by a warm PBS wash and incubated with Met-free DMEM (20 min) before addition of inhibitors (5 – 10 min) to block either cytosolic (50 μg/ml emetine) or mitochondrial (100 μg/ml chloramphenicol) protein synthesis. HPG was then added (500 μM final concentration) and cells incubated for 2.3 hours under normal culture conditions, after which mitochondria were prepared as previously reported (46). Mitochondria were solubilised with 0.4% (w/v) SDS at 40 μg protein/60 μl in PBS for 10 min at room temperature, followed by centrifugation (20 k*g*, 5 min). The solubilised proteins in the supernatant were used for the click reaction in a final volume of 120 μl (0.2% w/v final SDS concentration) with the following components added in order to the indicated final concentrations: 20 μM picolyl-biotin-azide (Jena Bioscience CLK-1167-5), 1.2 mM BTTAA (Jena Bioscience, CLK-067-100), 600 μM CuSO_4_ and 5 mM sodium ascorbate. The click reaction was incubated for 60 min at 25° C and terminated by standard methanol/chloroform protein precipitation. Precipitated proteins were resolubilised in 1x Bis-Tris SDS-PAGE sample buffer and separated on 12% Bis-Tris acrylamide (29:1 Acryl:Bis) gels using 1x MOPS running buffer supplemented with 5 mM sodium bisulphite as a reducing agent. Proteins were transferred onto PVDF membranes, and blocked (5% BSA, TTBS) prior to incubation with Streptavidin-HRP and visualisation by ECL.

## Acknowledgments

This work was supported by The Wellcome Trust [203105/Z/16/Z to RNL and ZCL] and EU Horizon2020 MCS [ITN REMIX]. We are grateful to Rob Taylor and Gavin Falkous for providing the mS22 deficient patient fibroblast cell line.

## References

1. Palade GE (1952) The fine structure of mitochondria. Anat Rec 114(3):427–451.

2. Sjostrand FS (1953) Electron microscopy of mitochondria and cytoplasmic double membranes. Nature 171(4340):30–32.

3. Frey TG & Mannella CA (2000) The internal structure of mitochondria. Trends Biochem Sci 25(7):319–324.

4. Jakobs S, Stephan T, Ilgen P, & Bruser C (2020) Light Microscopy of Mitochondria at the Nanoscale. Annu Rev Biophys.

5. Glancy B (2020) Visualizing Mitochondrial Form and Function within the Cell. Trends Mol Med 26(1):58–70.

6. Chacinska A, et al. (2003) Mitochondrial translocation contact sites: separation of dynamic and stabilizing elements in formation of a TOM-TIM-preprotein supercomplex. The EMBO journal 22(20):5370–5381.

7. Donzeau M, et al. (2000) Tim23 links the inner and outer mitochondrial membranes. Cell 101(4):401–412.

8. Gilkerson RW, Selker JM, & Capaldi RA (2003) The cristal membrane of mitochondria is the principal site of oxidative phosphorylation. FEBS Lett 546(2-3):355–358.

9. Vogel F, Bornhovd C, Neupert W, & Reichert AS (2006) Dynamic subcompartmentalization of the mitochondrial inner membrane. The Journal of cell biology 175(2):237–247.

10. Wilkens V, Kohl W, & Busch K (2013) Restricted diffusion of OXPHOS complexes in dynamic mitochondria delays their exchange between cristae and engenders a transitory mosaic distribution. Journal of cell science 126(Pt 1):103–116.

11. Strauss M, Hofhaus G, Schroder RR, & Kuhlbrandt W (2008) Dimer ribbons of ATP synthase shape the inner mitochondrial membrane. The EMBO journal 27(7):1154–1160.

12. Harner M, et al. (2011) The mitochondrial contact site complex, a determinant of mitochondrial architecture. The EMBO journal 30(21):4356–4370.

13. Pfanner N, et al. (2014) Uniform nomenclature for the mitochondrial contact site and cristae organizing system. The Journal of cell biology 204(7):1083–1086.

14. Rabl R, et al. (2009) Formation of cristae and crista junctions in mitochondria depends on antagonism between Fcj1 and Su e/g. The Journal of cell biology 185(6):1047–1063.

15. Bogenhagen DF, Rousseau D, & Burke S (2008) The layered structure of human mitochondrial DNA nucleoids. The Journal of biological chemistry 283(6):3665–3675.

16. Brown TA, et al. (2011) Superresolution fluorescence imaging of mitochondrial nucleoids reveals their spatial range, limits, and membrane interaction. Mol Cell Biol 31(24):4994–5010.

17. Kukat C, et al. (2011) Super-resolution microscopy reveals that mammalian mitochondrial nucleoids have a uniform size and frequently contain a single copy of mtDNA. Proceedings of the National Academy of Sciences of the United States of America 108(33):13534–13539.

18. Satoh M & Kuroiwa T (1991) Organization of multiple nucleoids and DNA molecules in mitochondria of a human cell. Experimental cell research 196(1):137–140.

19. Iborra FJ, Kimura H, & Cook PR (2004) The functional organization of mitochondrial genomes in human cells. BMC Biol 2:9.

20. Antonicka H, Sasarman F, Nishimura T, Paupe V, & Shoubridge EA (2013) The mitochondrial RNA-binding protein GRSF1 localizes to RNA granules and is required for posttranscriptional mitochondrial gene expression. Cell metabolism 17(3):386–398.

21. Jourdain AA, et al. (2013) GRSF1 regulates RNA processing in mitochondrial RNA granules. Cell metabolism 17(3):399–410.

22. Antonicka H & Shoubridge EA (2015) Mitochondrial RNA Granules Are Centers for Posttranscriptional RNA Processing and Ribosome Biogenesis. Cell reports 10(6):920–932.

23. Tu YT & Barrientos A (2015) The Human Mitochondrial DEAD-Box Protein DDX28 Resides in RNA Granules and Functions in Mitoribosome Assembly. Cell reports 10(6):854–864.

24. Stoldt S, et al. (2018) Spatial orchestration of mitochondrial translation and OXPHOS complex assembly. Nature cell biology 20(5):528–534.

25. Herrmann JM, Woellhaf MW, & Bonnefoy N (2013) Control of protein synthesis in yeast mitochondria: the concept of translational activators. Biochimica et biophysica acta 1833(2):286–294.

26. Weraarpachai W, et al. (2009) Mutation in TACO1, encoding a translational activator of COX I, results in cytochrome c oxidase deficiency and late-onset Leigh syndrome. Nature genetics 41(7):833–837.

27. Estell C, Stamatidou E, El-Messeiry S, & Hamilton A (2017) In situ imaging of mitochondrial translation shows weak correlation with nucleoid DNA intensity and no suppression during mitosis. Journal of cell science 130(24):4193–4199.

28. Kolb HC, Finn MG, & Sharpless KB (2001) Click Chemistry: Diverse Chemical Function from a Few Good Reactions. Angewandte Chemie 40(11):2004–2021.

29. Dieterich DC, et al. (2010) In situ visualization and dynamics of newly synthesized proteins in rat hippocampal neurons. Nature neuroscience 13(7):897–905.

30. Hinz FI, Dieterich DC, Tirrell DA, & Schuman EM (2012) Non-canonical amino acid labeling in vivo to visualize and affinity purify newly synthesized proteins in larval zebrafish. ACS chemical neuroscience 3(1):40–49.

31. Zorkau M, et al. (2020) Visualizing mitochondrial ribosomal RNA and mitochondrial protein synthesis in human cell lines.. Meths Mol. Biol. In Press.

32. Rostovtsev VV, Green LG, Fokin VV, & Sharpless KB (2002) A stepwise huisgen cycloaddition process: copper(I)-catalyzed regioselective “ligation” of azides and terminal alkynes. Angewandte Chemie 41(14):2596–2599.

33. Spencer AC & Spremulli LL (2004) Interaction of mitochondrial initiation factor 2 with mitochondrial fMet-tRNA. Nucleic acids research 32(18):5464–5470.

34. Arguello T, Kohrer C, RajBhandary UL, & Moraes CT (2018) Mitochondrial methionyl N-formylation affects steady-state levels of oxidative phosphorylation complexes and their organization into supercomplexes. The Journal of biological chemistry 293(39):15021–15032.

35. Chomyn A (1996) In vivo labeling and analysis of human mitochondrial translation products. Methods in enzymology 264:197–211.

36. Bogenhagen DF & Haley JD (2020) Pulse-chase SILAC-based analyses reveal selective oversynthesis and rapid turnover of mitochondrial protein components of respiratory complexes. The Journal of biological chemistry 295(9):2544–2554.

37. Wesolowska M, et al. (2015) Adult Onset Leigh Syndrome in the Intensive Care Setting: A Novel Presentation of a C12orf65 Related Mitochondrial Disease. Journal of neuromuscular diseases 2(4):409–419.

38. John GB, et al. (2005) The mitochondrial inner membrane protein mitofilin controls cristae morphology. Molecular biology of the cell 16(3):1543–1554.

39. Kehrein K, et al. (2015) Organization of Mitochondrial Gene Expression in Two Distinct Ribosome-Containing Assemblies. Cell reports 10(6):843–853.

40. Mannella CA, et al. (2001) Topology of the mitochondrial inner membrane: dynamics and bioenergetic implications. IUBMB life 52(3-5):93–100.

41. Mannella CA, Lederer WJ, & Jafri MS (2013) The connection between inner membrane topology and mitochondrial function. Journal of molecular and cellular cardiology 62:51–57.

42. Wolf DM, et al. (2019) Individual cristae within the same mitochondrion display different membrane potentials and are functionally independent. The EMBO journal 38(22):e101056.

43. Bogenhagen DF, Martin DW, & Koller A (2014) Initial steps in RNA processing and ribosome assembly occur at mitochondrial DNA nucleoids. Cell metabolism 19(4):618–629.

44. Stephan T, Roesch A, Riedel D, & Jakobs S (2019) Live-cell STED nanoscopy of mitochondrial cristae. Scientific reports 9(1):12419.

45. Wang C, et al. (2019) A photostable fluorescent marker for the superresolution live imaging of the dynamic structure of the mitochondrial cristae. Proceedings of the National Academy of Sciences of the United States of America 116(32):15817–15822.

46. Richter R, et al. (2010) A functional peptidyl-tRNA hydrolase, ICT1, has been recruited into the human mitochondrial ribosome. The EMBO journal 29(6):1116–1125.

