## Supplemental information for "High resolution imaging of nascent mitochondrial protein synthesis in cultured human cells"

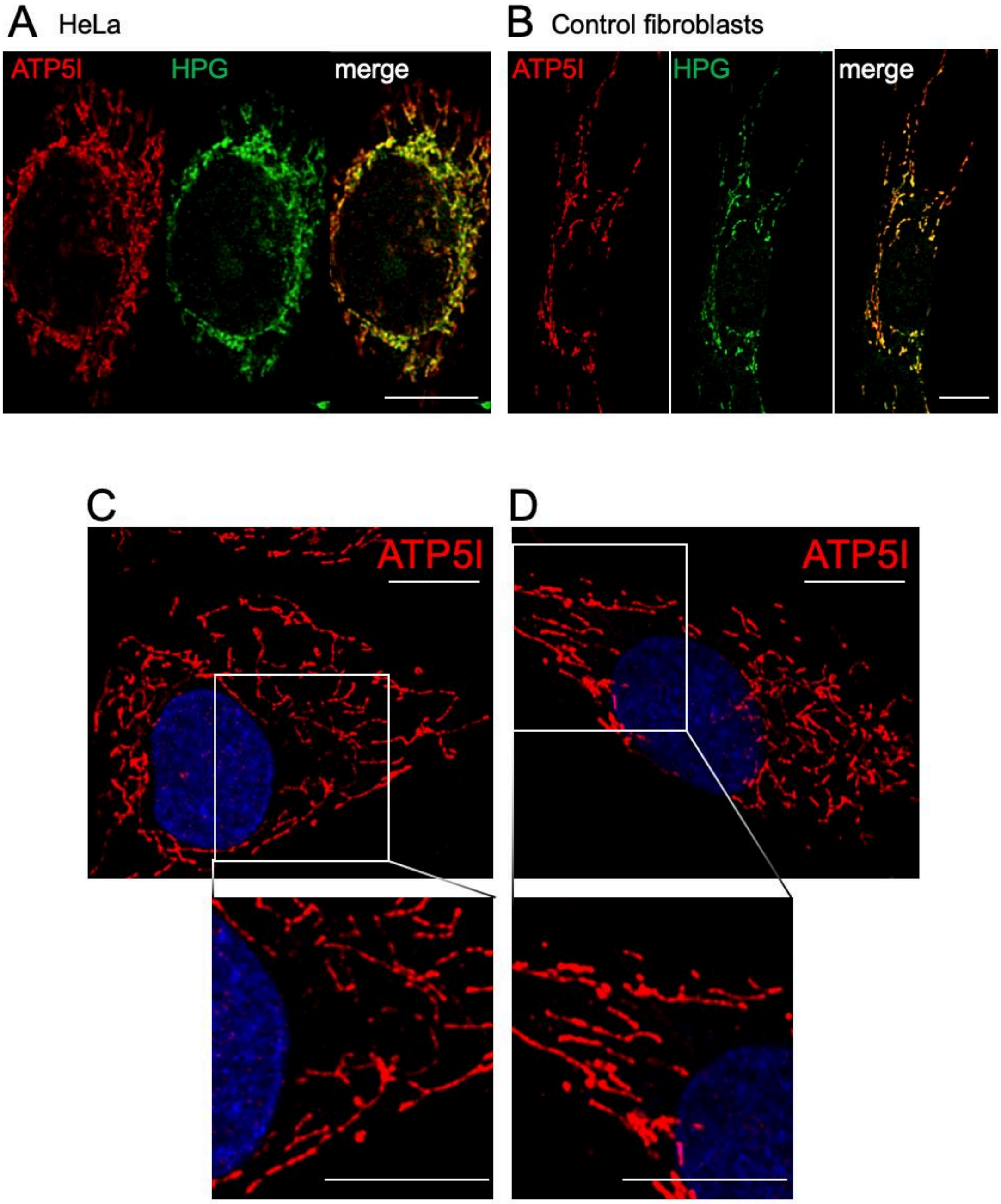


**Figure S1. Mitochondrial protein synthesis can be efficiently visualised within the mitochondrial reticulum in multiple cell types.**
HeLa cells (A) and control dermal fibroblasts (B) were pulsed with HPG for 25 mins in the presence of the cytosolic translation inhibitor cycloheximide, followed by the click reaction as described in Methods. Cells were then fixed and stained with antibodies against ATP5I. Representative deconvolved confocal images are shown. Scale bars = 10 μm.
**To determine whether the integrity of the mitochondria network was compromised due to the FUNCAT procedure,** control fibroblasts were left untreated (C), or subjected to an HPG incubation (15 mins, with cycloheximide) and brief digitonin permeabilisation (80 sec) to remove unincorporated HPG, followed by the standard click reaction as given in Methods (D)**.** Cells were fixed and stained for a marker of mitochondrial cristae (OXPHOS complex V, ATP5I). Representative deconvolved confocal images are shown, panels C and D show nuclei counterstained with Hoechst, and scale bars in all panels = 10 μm.

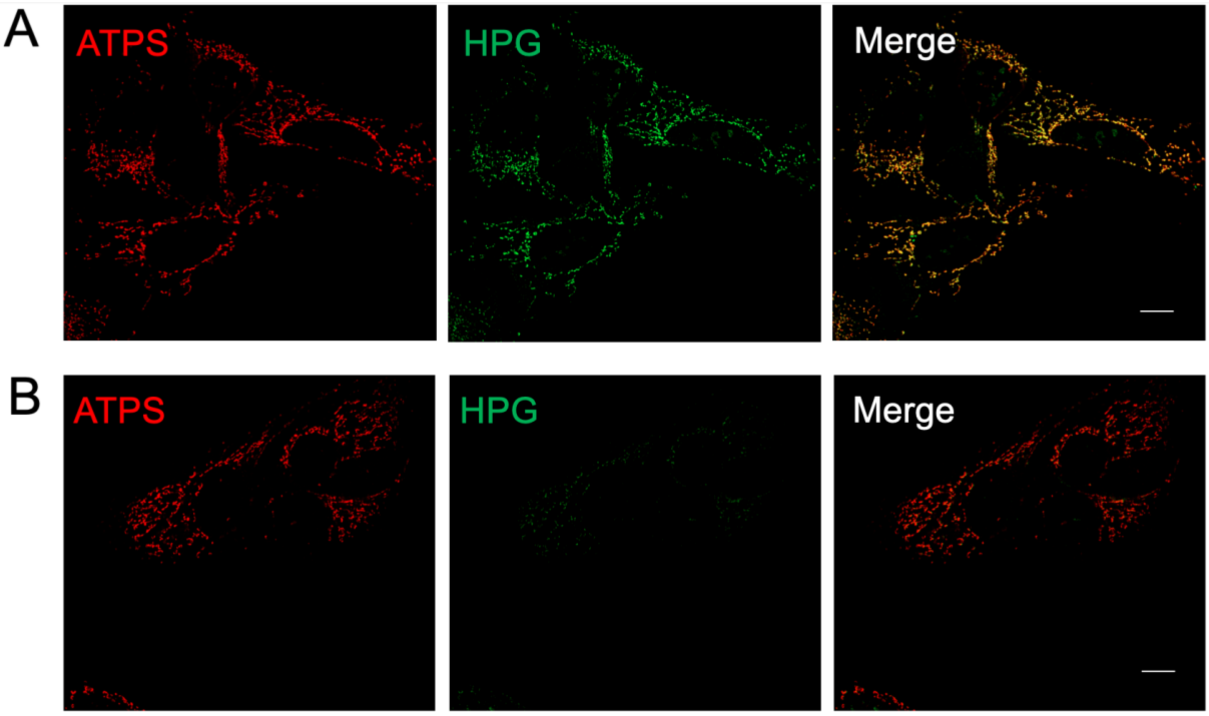


**Figure S2. To determine whether mitochondrial protein synthesis could initiate *de novo* with HPG as a methionine substitute, cells were pre-treated with puromycin.**
Control fibroblasts were pretreated with puromycin (2 hr) to terminate synthesis, prior to addition of HPG. Cells were then incubated with HPG (2 hr) in the absence (A) or continued presence of puromycin (B). Cycloheximide inhibition of cytosolic translation was maintained throughout. Post click reaction and fixation, cells were stained with antibodies against the Complex V protein ATP5B (ATPS) as a marker of the mitochondrial network. Representative deconvolved confocal images are shown with scale bars = 10 μm.

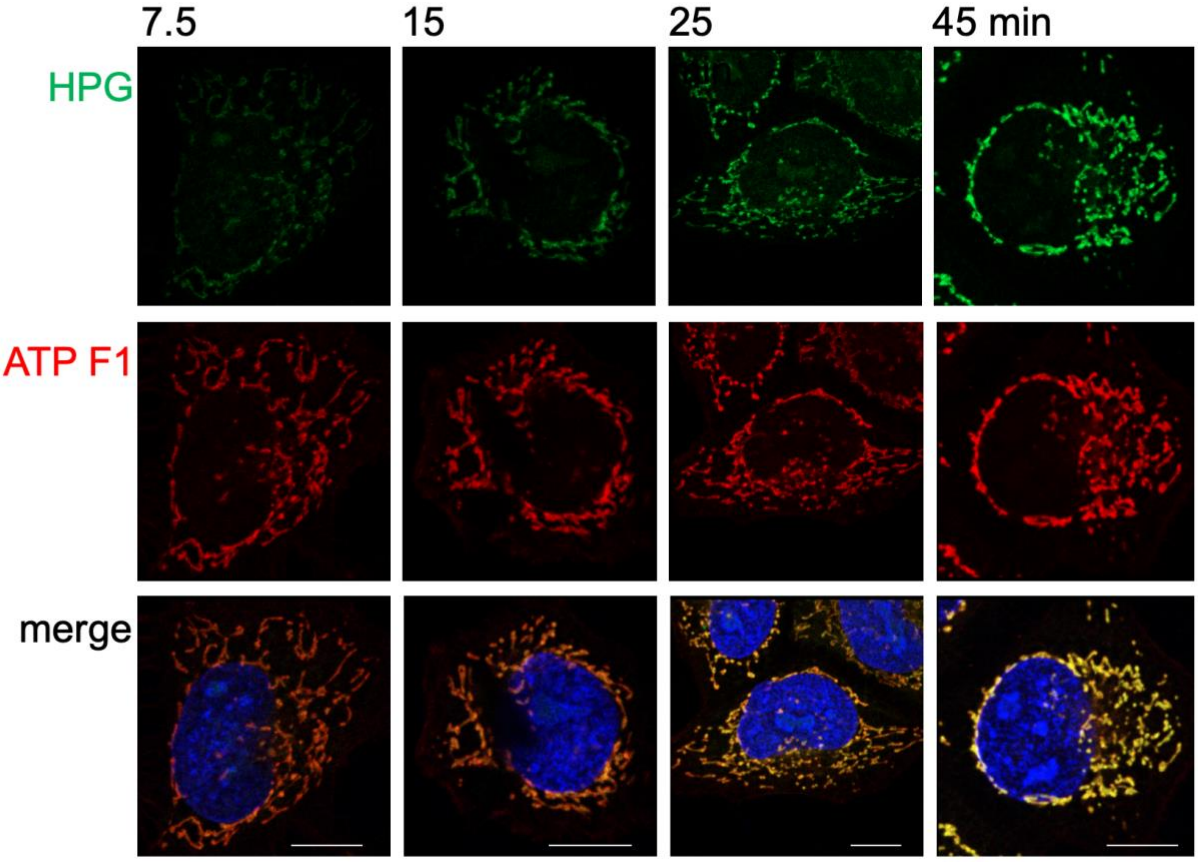


**Figure S3. Mitochondrial protein synthesis was monitored in HeLa cells.** HeLa cells were cultured in HPG (green) and cycloheximide for the pulse times indicated. Cells were then fixed and stained with antibodies against a member of Complex V (ATP F1, red) of the OXPHOS machinery to highlight the mitochondrial network. Nuclei are stained with Hoechst (blue) in the merged images. Representative confocal images are shown and all scale bars = 10 µm.

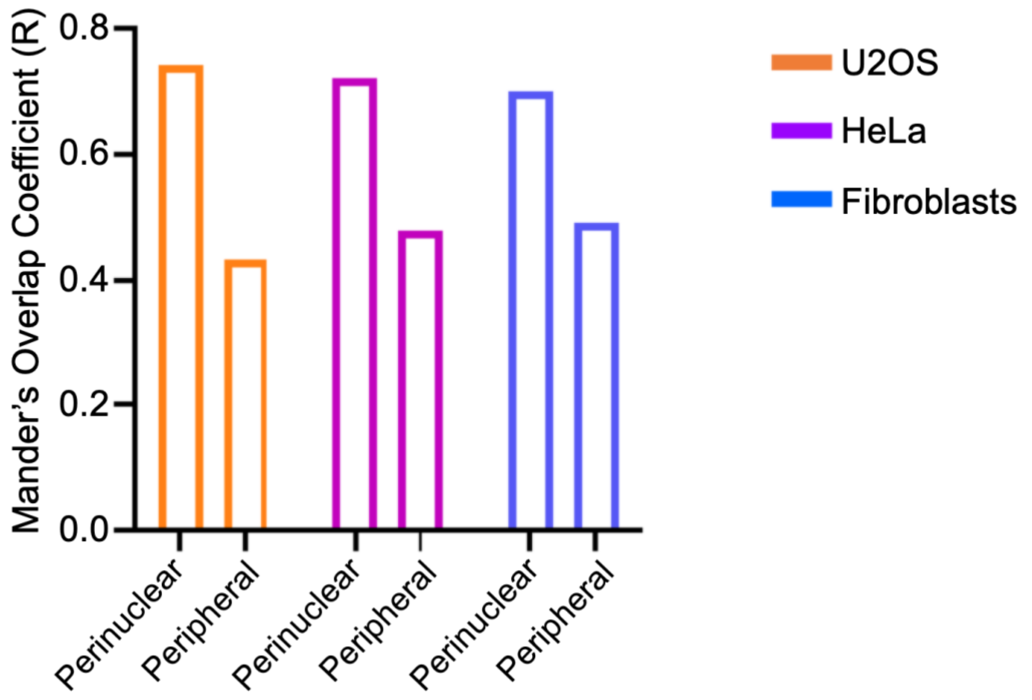


**Figure S4. Distribution of mitochondrial protein synthesis between the periphery and the perinuclear region remains similar in different cell types.** HPG labelling with subsequent immunostaining for ATP5I, as a marker of the mitochondrial network, was performed in U2OS, HeLa and fibroblast cells. Mander's overlap coefficient was calculated to reveal proportion of ATP5I covered by HPG in areas of the mitochondrial network in the perinuclear or peripheral regions of the cell.

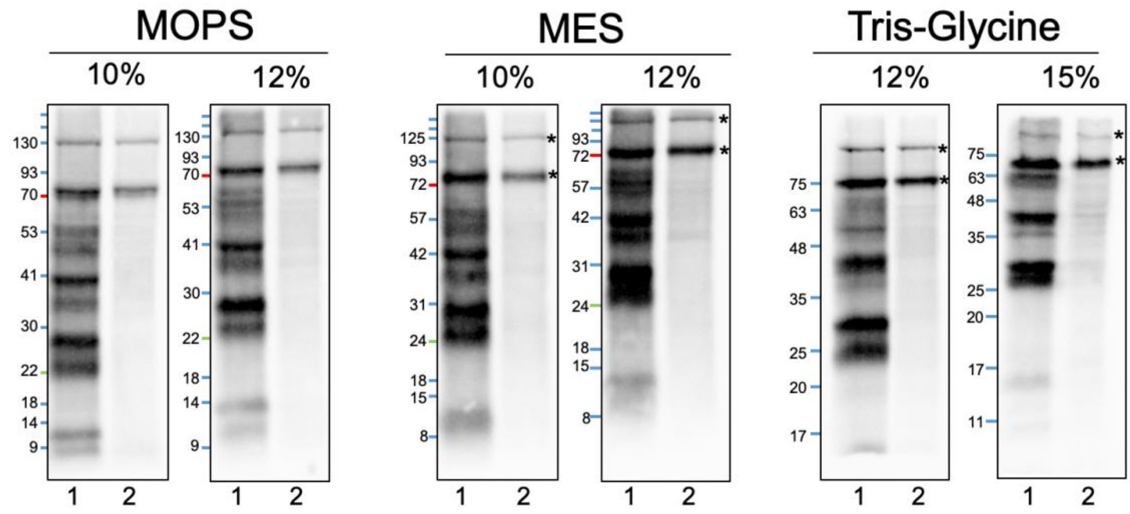


**Figure S5. Different conditions and reagent combinations were tested during optimisation of BONCAT signal of mitochondrial translation**

Uninduced Flp-in T-Rex 293 cells carrying a COX8-BioID2 cassette, were incubated with emetine to inhibit cytosolic translation and pulsed with HPG (500 mM, 2 h), in the absence (1), or presence of chloramphenicol (2) as a control that prevents mitochondrial translation. Mitochondria were isolated and samples 1 and 2 were solubilised with SDS (0.2%) and the click reaction performed with p-azide-biotin. Aliquots of samples 1 and 2 were electrophoresed through 6 variants of 29:1 acrylamide/ bisacrylamide gels. MOPS and MES buffers were used with Bis-Tris gels (10%, 12%), and Tris-Glycine running buffer was used with Tris-Glycine gels (12%, 15%). Following wet transfer, membranes were probed with streptavidin-HRP, and signals visualised with ECL. Asterisks indicate endogenously biotinylated proteins. Representative images are shown.

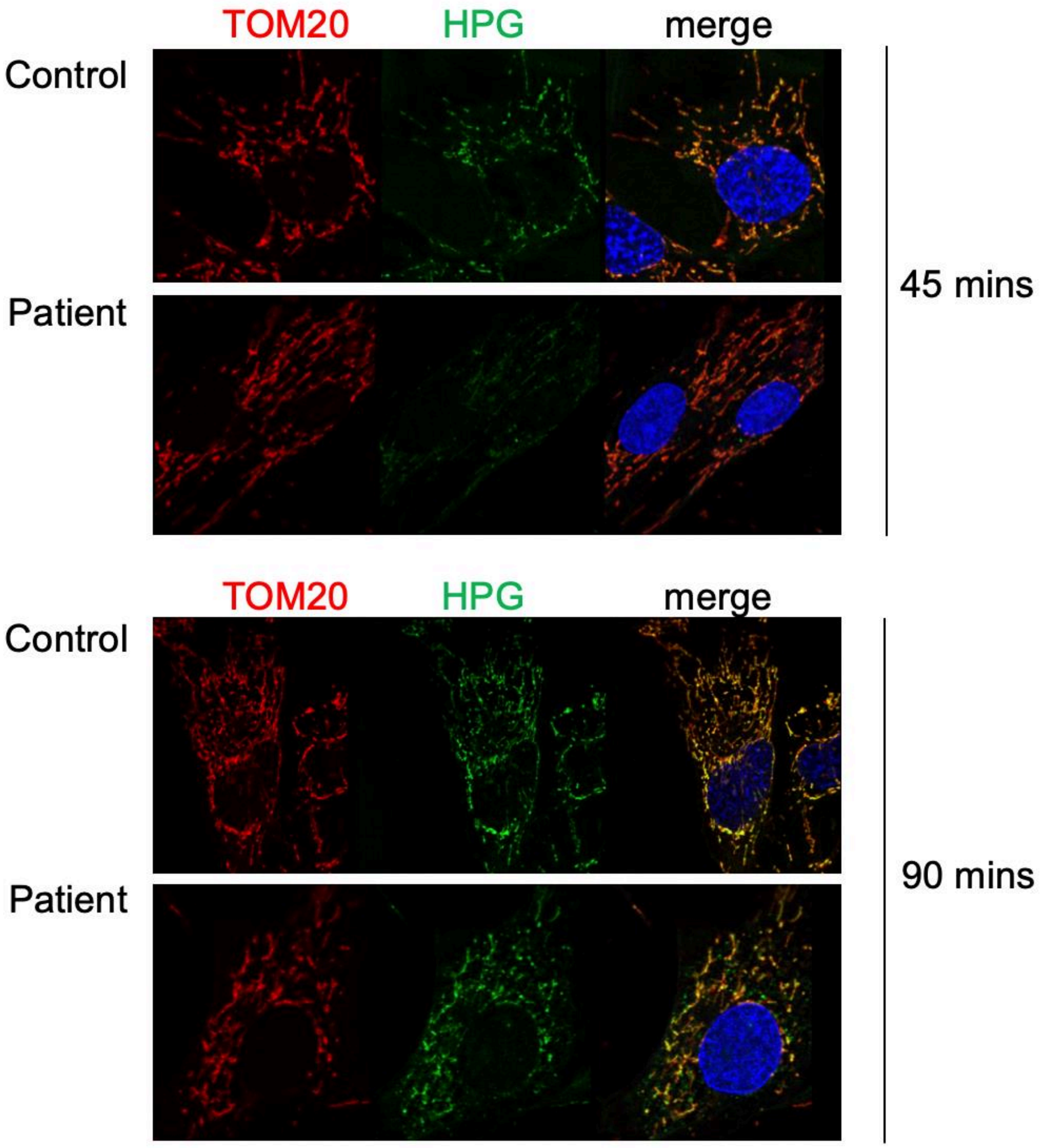


**Figure S6. Mitochondrial protein synthesis defects can be seen in patients with mitochondrial dysfunction.** Dermal fibroblasts from control and a patient with a mutation in C12orf65, known to cause a defect in mitochondrial translation, were were subjected to 45 and 90 minute HPG pulses. Cells were then fixed and stained with antibodies against an outer membrane marker (TOM20) to highlight the mitochondrial network. Nuclei are stained with Hoechst (blue) in the merged images. Representative confocal images are shown.

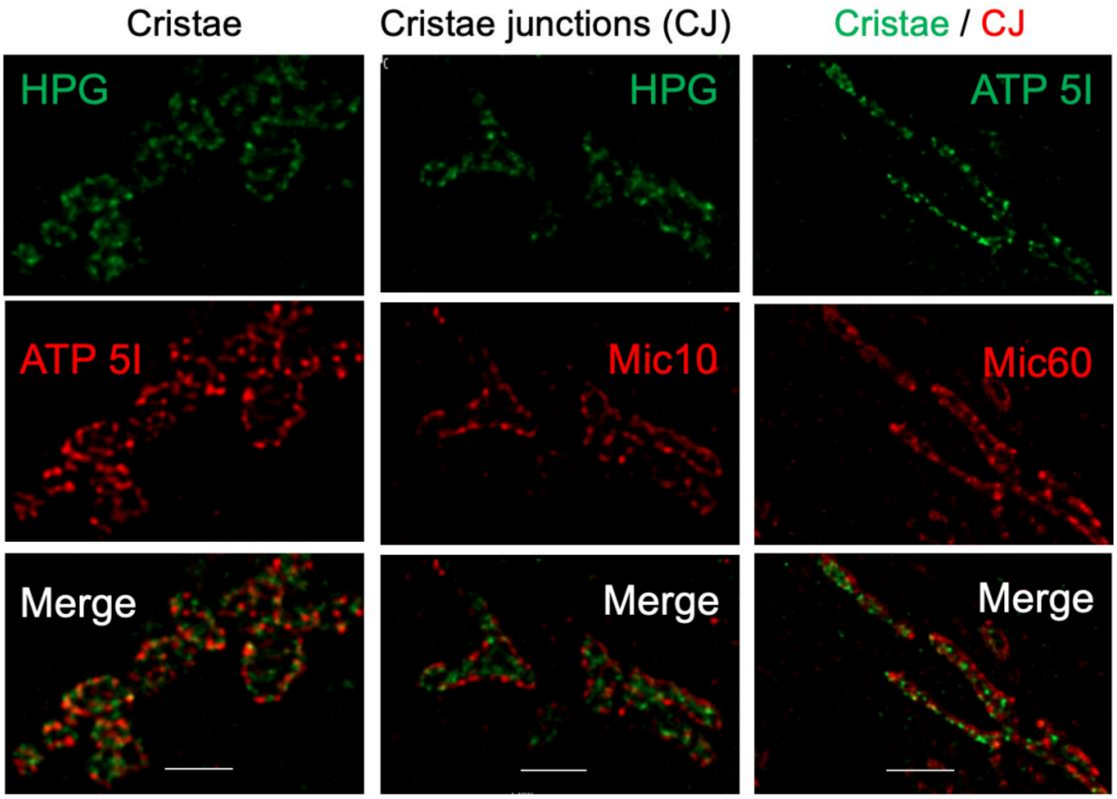


**Figure S7. Mitochondrial protein synthesis occurs at the cristae in preference to cristae junctions.** HeLa cells were pulsed with HPG (green) for 30 mins in the presence of cycloheximide followed by click reactions performed as described in Methods. Mitochondrial cristae (left) and cristae junctions (centre) were visualised (red) with antibodies against ATP5I and Mic10 respectively. To demonstrate the relative positions of cristae and cristae junctions, fixed cells were concurrently stained with antibodies against ATP5I and Mic60 respectively (right). Representative deconvolved super resolution images are shown, all scale bars = 1 μm.

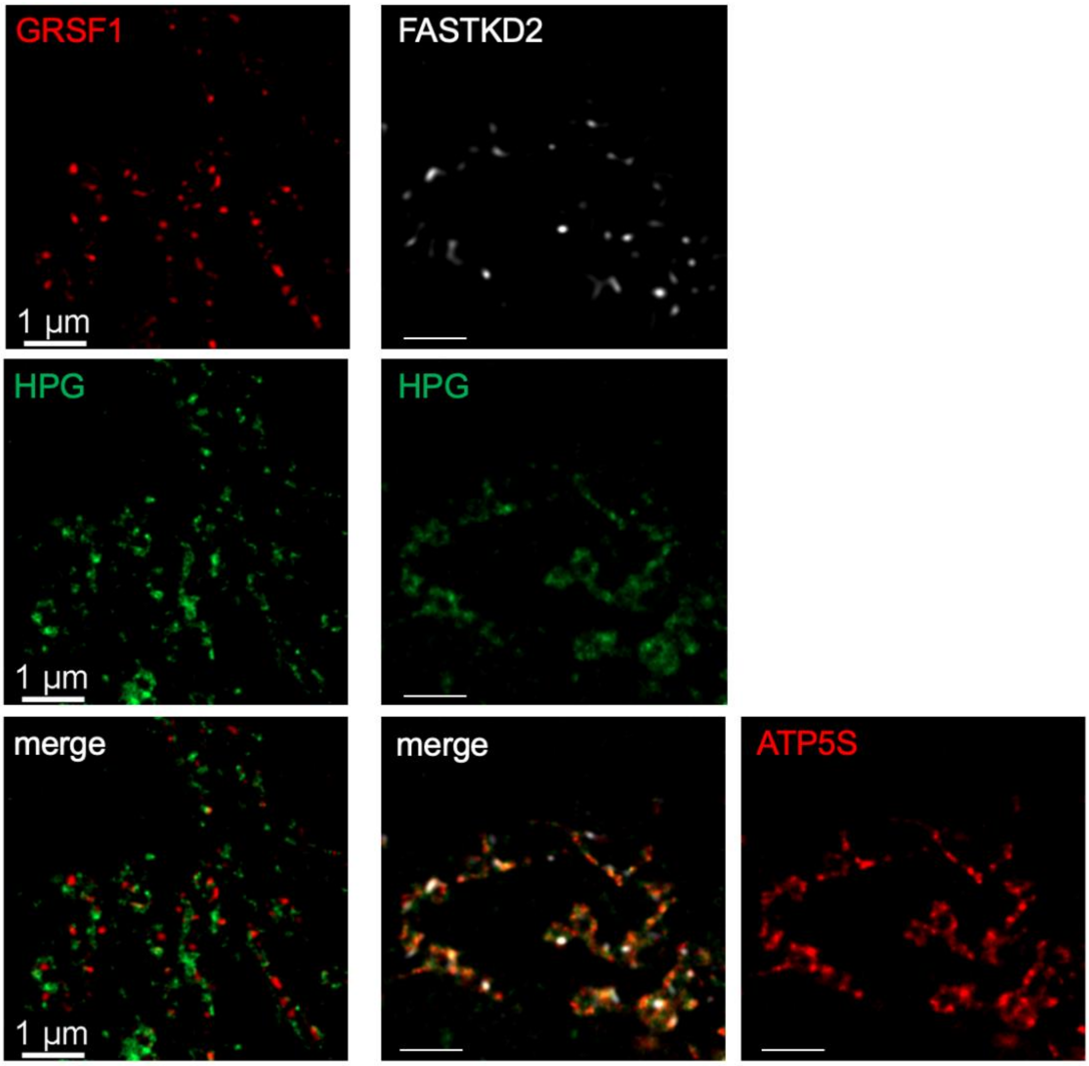


**Figure S8. Mitochondrial protein synthesis colocalises more closely with cristae than with mitochondrial RNA granules.** U2OS were subjected to a short pulse (7.5 min) of HPG (green) in the presence of cycloheximide, followed by immunostaining as described in Methods. GRSF1 (red) and FASTKD2 (white) were used independently as markers to determine the location of mitochondrial RNA granules. In addition to HPG and FASTKD2, cristae were also decorated with antibodies against the beta subunit of complex V (ATP5S) and the merged image is shown. Representative deconvolved confocal images are shown, all scale bars = 1 µm.

Table S1. Primary antibodies used in this study

| Antigen | Dilution | Host species and isotype | Supplier | Catalogue number |
|---|---|---|---|---|
| TOM20 | 1:400 | Rabbit IgG | Abcam | ab78547 |
| TIM23 | 1:200 | Rabbit IgG | Protein Tech | 11123-1-AP |
| Mic60 | 1:400 | Mouse IgG | Abcam | ab110329 |
| Mic10 | 1:200 | Rabbit IgG | Abcam | ab84969 |
| MTCO1 | 1:200 | Mouse IgG | Abcam | ab14705 |
| ATP5I | 1:200 | Rabbit IgG | Protein Tech | 16483-1-AP |
| ATPS | 1:200 | Mouse IgG | Abcam | ab14730 |
| FASTKD2 | 1:200 | Rabbit IgG | Protein Tech | 17464-1-AP |
| GRSF1 | 1:200 | Rabbit IgG | Abcam | ab194358 |
| MRPS15 | 1:200 | Rabbit IgG | Protein Tech | 17006-1-AP |
